## Supplemental for "Deep Template-based Protein Structure Prediction"

### Supplementary information for Deep Template-based Protein Structure Prediction

Fandi Wu[1,2,3], Jinbo Xu[1,*]

[1]Toyota Technological Institute at Chicago, Chicago, IL 60637, USA, [2]Institute of Computing Technology, Chinese Academy of Sciences, Beijing, 626011, China and [3]University of Chinese Academy of Sciences, Beijing, 100049, China

#### Representation of protein alignment

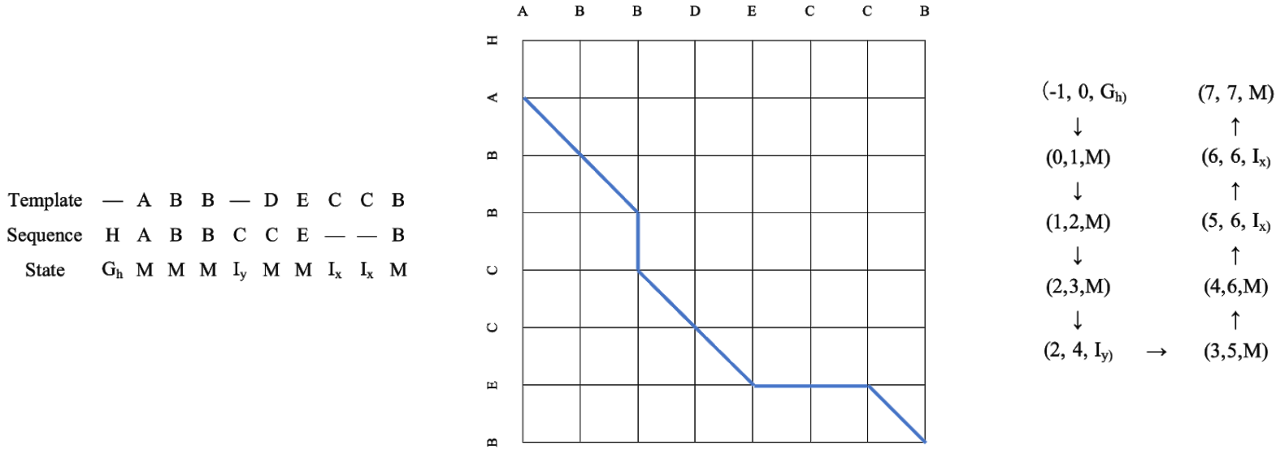


Figure S1. Three different representations of a protein alignment. Left: a sequence of alignment states; Middle: a path in the alignment matrix; Right: a sequence of triples, each composed of two residue indices and one alignment state.

#### State transition matrix for DRNF method

Table S1. The state transition score matrix for DRNF (deep convolutional residual neural fields), in which an entry of -inf indicates a forbidden transition. M denotes that two residues are aligned; $I_x$ and $I_y$ denote insertions at proteins x and y, respectively; and $G_h$ and $G_t$ denote head and tail gaps.

| | M | $I_x$ | $I_y$ | $G_h$ | $G_t$ |
|---|---|---|---|---|---|
| M | 0.5 | -5 | -5 | -inf | 0 |
| $I_x$ | 0 | -1 | -2 | -inf | -inf |
| $I_y$ | 0 | -inf | -1 | -inf | -inf |
| $G_h$ | 0 | -inf | -inf | 0 | -inf |
| $G_t$ | -inf | -inf | -inf | -inf | 0 |

### DRNF for protein alignment without distance information

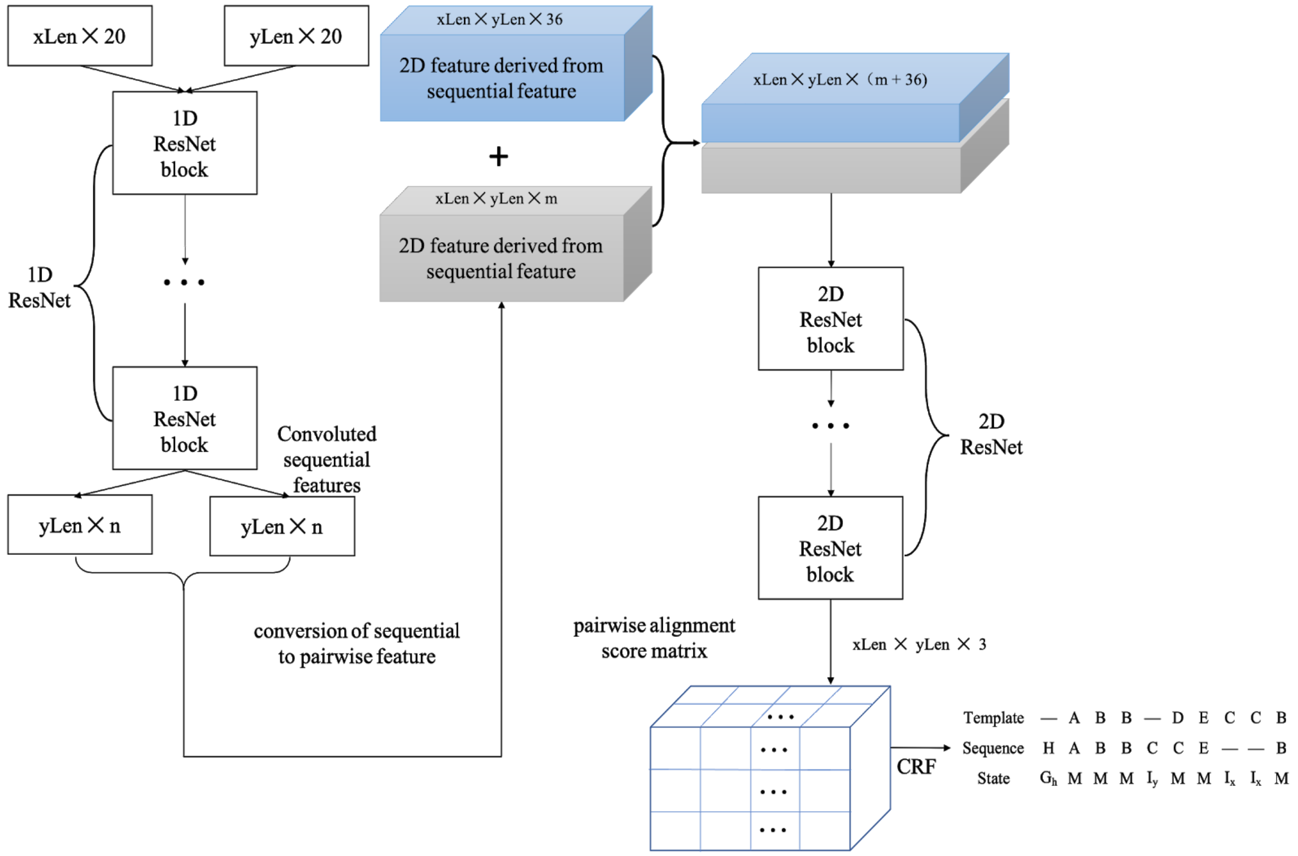


Figure S2. The deep network architecture for protein alignment without distance information.

### Protein alignment with distance information

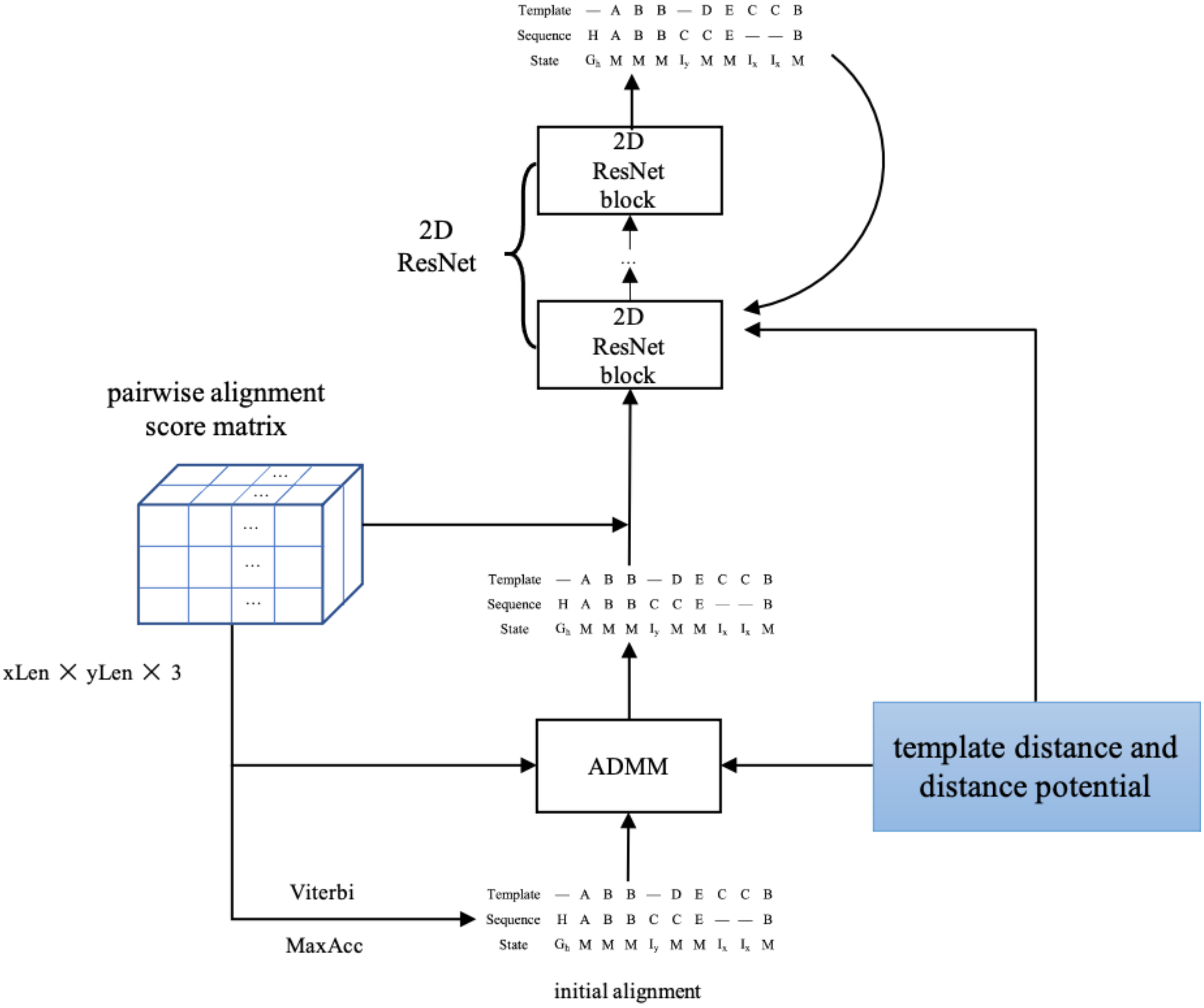


Figure S3. The deep network architecture for protein alignment with distance information.

### ADMM for protein alignment with predicted distance potential

We formulate the sequence-template alignment problem as the following integer quadratic programming problem.

$$max_z \; w \times \sum_{(i,j,u)\in Z} \theta_{ij}^u z_{ij}^u + \sum_{(i,j,u)\in Z,\,(k,l,v)\in Z} \theta_{ijkl}^{uv} z_{ij}^u z_{kl}^v \quad s.t. \; \sum_{j,u} z_{ij}^u = 1 \; for \; any \; i \tag{1}$$

where w is a weight factor with a default value 1. $z_{ij}^u$ is a binary variable that equals to 1 if and only if the triple (i, j, u) is in the alignment (see the representation of an alignment). $\theta_{ij}^u$ represents the score generated by DRNF for residues i and j with state u. $\theta_{ijkl}^{uv}$ is equal to 0 if either u or v is not the match state. Otherwise, it equals the potential of query residues j and l falling into a distance bin *d* where *d* is the distance bin into which the two template residues i and k fall.

It is computationally hard to optimize the score function in Eq.1 when gaps are allowed in the alignment. To use ADMM (Alternating Direction Method of Multipliers) algorithm described in (Ma, et al., 2014), we make a copy of z and add the constraint $z_{i,j}^u = y_{i,j}^u$ . Then, we add a term to penalize the difference between z and y. Eq.1 becomes a new quadratic problem.

$$Max_{\{z,y\}} \frac{w}{2} \sum_{(i,j,u)\in Z} \theta_{ij}^{u}(z_{ij}^{u} + y_{ij}^{u}) + \sum_{(i,j,u)\in Z,(k,l,v)\in Z} \theta_{ijkl}^{uv} z_{ij}^{u} y_{kl}^{v} - \frac{\rho}{2} \times \sum_{i,j,u} \left(z_{ij}^{u} - y_{kl}^{v}\right)^{2} \tag{2}$$

Where $\rho$ is a constant, and $y$ is a copy of $z$.

By using a Lagrange multiplier $\lambda$ for the constraints, we have the following Lagrange dual problem.

$$\begin{aligned} Max_{\{z,y\}} \frac{w}{2} \sum_{(i,j,u)\in Z} \theta_{ij}^{u}(z_{ij}^{u} + y_{ij}^{u}) + \sum_{(i,j,u)\in Z,(k,l,v)\in Z} \theta_{ijkl}^{uv} z_{ij}^{u} y_{kl}^{v} + \\ \sum_{i,j,u} \lambda_{ij}^{u} \left(z_{ij}^{u} - y_{kl}^{v}\right) - \frac{\rho}{2} \times \sum_{i,j,u} \left(z_{ij}^{u} - y_{kl}^{v}\right)^{2} \end{aligned} \tag{3}$$

Both z and y are binary variables, the last term in Eq. 3 can be expanded as follows.

$$\frac{\rho}{2} \times \sum_{i,j,u} \left(z_{ij}^{u} - z_{kl}^{v}\right)^{2} = \frac{\rho}{2} \sum_{i,j,u} \left(z_{ij}^{u} + z_{kl}^{v} - 2\, z_{ij}^{u} z_{kl}^{v}\right) \tag{4}$$

For a fixed $\lambda$, we can split Eq. 3 into the following two sub-problems,

$$y^{*} = argmax \sum_{k,l,v} y_{kl}^{v} C_{kl}^{v} \tag{5}$$

Where $C_{kl}^{v} = \frac{w}{2} \sum_{i,j,u} \theta_{kl}^{v} z_{ij}^{u} + \sum_{i,j,u} \theta_{ijkl}^{uv} z_{ij}^{u} - \lambda_{kl}^{v} - \frac{\rho}{2}\left(1 - 2z_{kl}^{v}\right)$

$$z^{*} = argmax \sum_{i,j,u} z_{ij}^{u} D_{ij}^{u} \tag{6}$$

Where $D_{ij}^{u} = \frac{w}{2} \sum_{k,l,v} \theta_{ij}^{u} z_{kl}^{v} + \sum_{k,l,v} \theta_{ijkl}^{uv} y_{kl}^{v} - \lambda_{ij}^{u} - \frac{\rho}{2}\left(1 - 2y_{ij}^{u}\right)$

Eq. 5 optimizes the objective function with respect to y while fixing z, Eq. 6 optimizes the objective function with respect to z while fixing y. Neither Eq. 5 or Eq. 6 has a quadratic term, so we can use the Viterbi algorithm to solve these two sub-problems. Briefly, the whole algorithm has the following main steps:

1. Run DRNF to generate an initial sequence-template alignment without using distance potential and apply this alignment to initialize z.
2. Fixing z, run the Viterbi algorithm to maximize Eq. 5 and update y. This will generate a new alignment specified by y.
3. Fix y and similarly run the Viterbi algorithm to maximize Eq. 6 and update z. This will generate a new alignment specified by z.
4. Check and update the best alignment using z according to the selection score.
5. If z and y are very close to each other, stop and yield the best alignment. Otherwise, update Lagrange multiplier $\lambda$ by

$$\lambda^{n+1} = \lambda^{n} - \rho\,(z^{*} - y^{*})$$

and repeat steps 2 and 3.

### Training analysis of DRNF for protein alignment

We have trained 4 different DRNF models by maximum-likelihood and show their learning curves in Fig. S1. The training and validation loss is calculated as the average negative log-likelihood of the reference alignments. Model1 and Model3 use secondary structure and solvent accessibility, so they have lower training and validation loss than Model2 and Model4, respectively. Model3 and Model4 use the TMalign alignments as references while Model1 and Model2 use the DeepAlign alignments as references. Model3 and Model4 have larger training and validation loss than Model1 and Model2. This is because TMalign aligns two protein structures using only geometric similarity while DeepAlign also takes into consideration sequence evolutionary information. That is, the DeepAlign alignments can be learned more easily by DRNF since both DeepAlign and DRNF make use of evolutionary information. During training, we pack several short protein pairs into a single minibatch. But in validation, we treat each protein pair as a single minibatch. Short protein pairs tend to have lower loss because of a smaller search space for alignments. Therefore, the average training loss in an epoch usually is larger than the average validation loss of the same epoch.

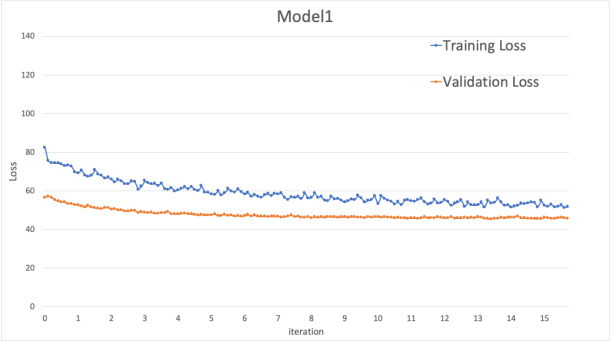


(a) Model1 training and validation loss

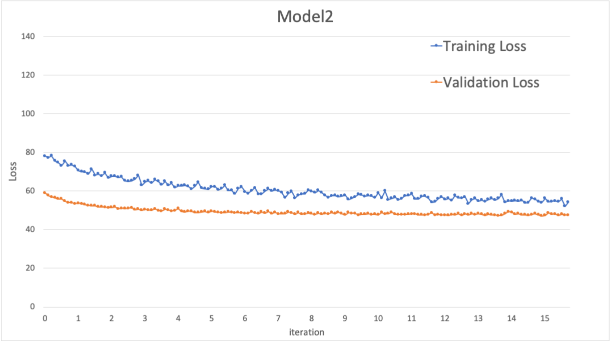


(b) Model2 training and validation loss

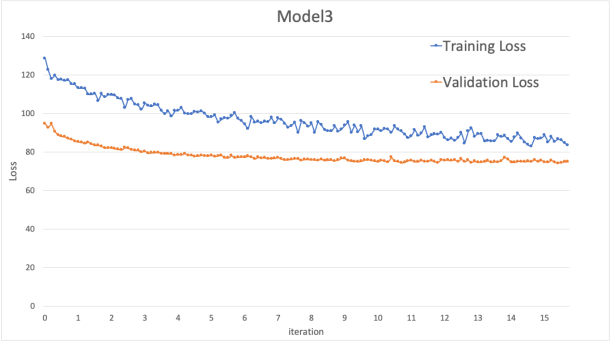


(c) Model3 training and validation loss

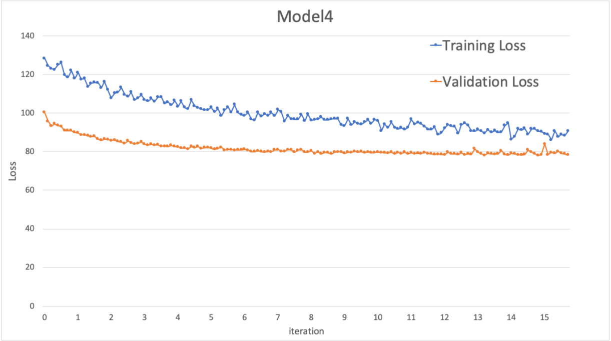


(d) Model4 training and validation loss

Figure S4. Training and validation loss of 4 different DRNF models. (a) Model1 is trained on DeepAlign alignments and uses structure information as features. (b) Model2 is trained on DeepAlign alignments but does not use structure information. (c) Model3 is trained on TMalign alignments and uses structure information as features. (d) Model4 is trained on TMalign alignments and uses structure information.

### Contribution of protein features and impact of reference alignments

We have tested our DRNF (Deep Convolutional Residual Neural Fields) method for query-template alignment in 6 different settings, as described in Table S2. These 6 settings include if (predicted) structure information is used for (query) template or not, how the reference alignments are generated and how the DRNF alignments are generated. Table S3 shows the reference-independent alignment accuracy of DRNF in these 6 settings. Model1 does not use any predicted structure features, and has the worst alignment quality among these 6 settings. Model2 uses 3-class secondary structure, 8-class secondary structure and solvent accessibility as features and improves the TM-score and GDT by 0.025 and 0.019 over Model1. Both Model1 and Model2 are trained by the reference alignments generated by DeepAlign. Model3 uses reference alignments generated by TMalign and performs slightly worse than Model2 when the sequence-template structure similarity falls into (0.8, 1]. Model4 is trained by the DeepAlign alignments and generates alignment by maximizing the expected accuracy (i.e., MaxAcc). On average Model4 slightly outperforms Model2 (which uses the Viterbi method to generate alignments) by 0.008 in terms of TMscore. However, Model4 slightly underperforms Model2 when the sequence-template similarity falls into (0.8, 1]. Further, the MaxAcc method takes a longer time than the Viterbi method to generate an alignment, so we prefer to use the Viterbi method to generate alignments. Models 5 and 6 use a deeper ResNet and achieve slightly better accuracy than Models 2 and 4, respectively.

Table S2. The setting of different DRNF method for query-template alignment

| | Structure Feature | Training data | Alignment generation | Layer |
|---|---|---|---|---|
| Model1 | × | DeepAlign | Viterbi | 20 |

| | | | | |
|---|---|---|---|---|
| Model2 | √ | DeepAlign | Viterbi | 20 |
| Model3 | √ | TMalign | Viterbi | 20 |
| Model4 | √ | DeepAlign | MaxAcc | 20 |
| Model5 | √ | DeepAlign | Viterbi | 50 |
| Model6 | √ | DeepAlign | MaxAcc | 50 |

Table S3. The contribution of protein features and impact of reference alignments. GDT is rescaled to [0, 1].

| | DRNF_Model1 | | DRNF_Model2 | | DRNF_Model3 | | DRNF_Model4 | | DRNF_Model5 | | DRNF_Model6 | |
|---|---|---|---|---|---|---|---|---|---|---|---|---|
| | TMscore | GDT | TMscore | GDT | TMscore | GDT | TMscore | GDT | TMscore | GDT | TMscore | GDT |
| (0,1] | 0.550 | 0.413 | 0.525 | 0.432 | 0.526 | 0.430 | 0.534 | 0.435 | 0.527 | 0.434 | 0.539 | 0.440 |
| (0.45, 0.55] | 0.345 | 0.270 | 0.380 | 0.294 | 0.386 | 0.296 | 0.391 | 0.298 | 0.379 | 0.295 | 0.397 | 0.303 |
| (0.55, 0.65] | 0.471 | 0.373 | 0.439 | 0.389 | 0.497 | 0.390 | 0.510 | 0.398 | 0.490 | 0.387 | 0.506 | 0.396 |
| (0.65, 0.8] | 0.585 | 0.494 | 0.610 | 0.515 | 0.608 | 0.512 | 0.611 | 0.513 | 0.617 | 0.522 | 0.624 | 0.526 |
| (0.8, 1] | 0.709 | 0.636 | 0.723 | 0.648 | 0.709 | 0.634 | 0.719 | 0.643 | 0.722 | 0.648 | 0.724 | 0.648 |

### Relationship between alignment score and model quality

To evaluate the relationship between alignment quality and alignment score (when distance information is not used), we normalize the alignment score by the minimum of query and template lengths. Here in calculating TMscore, we use the smaller of the template length and query length as the normalization constant. Note in other sections we always use the query length as the normalization constant in calculating TMscore and GDT. Fig. S5 shows that there is a certain correlation between the normalized alignment score and TMscore of the models derived from the alignments generated by DRNF. The correlation coefficient is 0.537 and the trendline $R^2$ is 0.288. When the normalized alignment score >0.3, 76.2% of DRNF alignments have TMscore>0.5; When the normalized score is >0.4, 86.2% of DRNF alignments have TMscore>0.5.

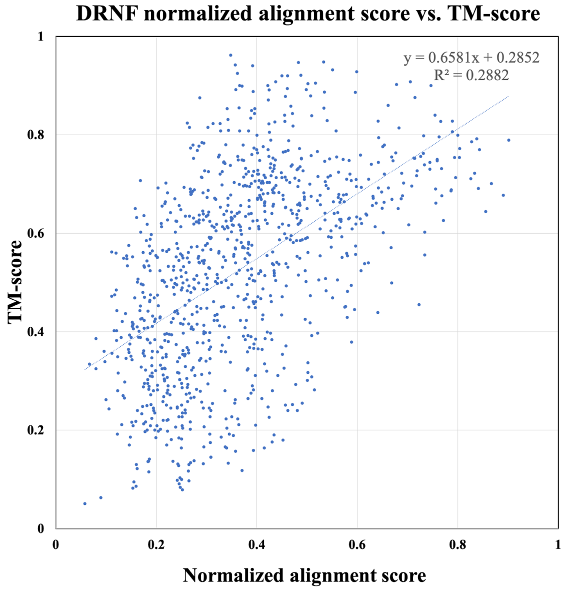


Figure S5. The relationship between the normalized alignment score and the alignment quality measured by TMscore.

### Comparison between NDThreader and structure alignment

We use TMalign to search the whole template database to find structurally the most similar templates for a CASP13 test target and then examine the difference between the NDThreader alignments and structure alignments. We use the target length as the normalization constant in calculating TMscore of a structure alignment. Fig. S6 shows the head-to-head comparison between the quality (TMscore) of the alignments generated by NDThreader and the TMscore of the structure alignments generated by TMalign. On FM, FM/TBM, TBM-hard, TBM-easy targets, the average TMscore of the most structurally similar templates is 0.584, 0.653, 0.764 and 0.844 while the 3D models built by MODELLER from NDThreader alignments have average TMscore 0.437, 0.578, 0.716 and 0.819, respectively, when the first-ranked templates are used. When the best of top 5 templates are used, NDThreader has average TMscore 0.473, 0.604, 0.743 and 0.826. This result indicates that for most FM and FM/TBM targets, there is still a gap between NDThreader alignment and structure alignment, and that for TBM targets NDThreader alignment is slightly worse than structure alignment.

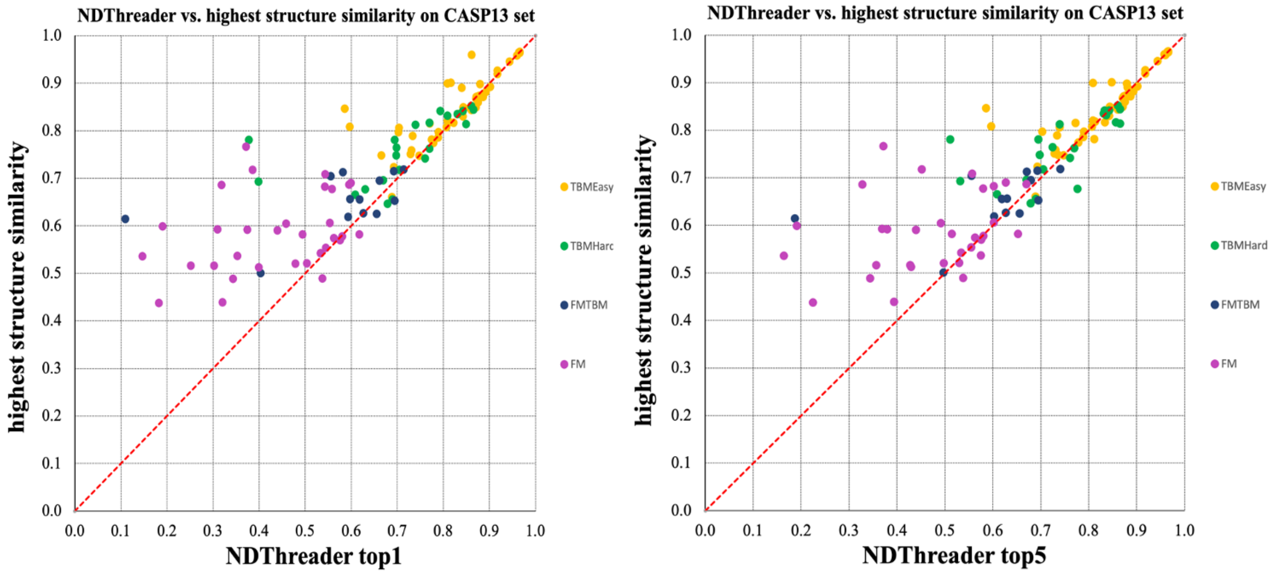


Figure S6. Head-to-head comparison between NDThreader alignments and TMalign alignments on the CASP13 targets. Left: top 1 models by NDThreader vs. TMalign. Right: the best of top 5 models by NDThreader vs. TMalign. Each point represents the quality (TM-score) of the model generated by NDThreader (x-axis) and the highest structure similarity returned by TMalign (y-axis).
